# Leptin Receptors in RIP-Cre^25Mgn^ neurons Mediate Anti-Dyslipidemia Effects of Leptin in Insulin-Deficient Male Mice

**DOI:** 10.1101/2020.03.13.989442

**Authors:** Ashish Singha, Juan Pablo Palavicini, Meixia Pan, Darleen Sandoval, Xianlin Han, Teppei Fujikawa

**Author notes:** Corresponding Author: Teppei Fujikawa.

## Abstract

Leptin is a potent endocrine hormone produced by adipose tissue and regulates a broad range of metabolism including glucose and lipid metabolism, with and without insulin. It is evident that central leptin signaling can lower hyperglycemia in insulin-deficient rodents via multiple mechanisms including restoration of dyslipidemia. However, the specific neurons that regulate these glucose-lowering and anti-dyslipidemia effects of leptin remain unidentified. Here we report that leptin receptors (LEPRs) in neurons expressing Cre recombinase driven by a short fragment of a promoter region of *Ins2* gene (RIP-Cre^25Mgn^ neurons) are required for central leptin signaling to reverse hyperglycemia and dyslipidemia in insulin-deficient mice. Ablation of LEPRs in RIP-Cre^25Mgn^ neurons completely blocks glucose-lowering effects of leptin in insulin-deficient mice. Further investigations reveal that insulin-deficient mice lacking LEPRs in RIP-Cre^25Mgn^ neurons (RIP-Cre^ΔLEPR^ mice) exhibit greater lipid levels in blood and liver compared to wild-type controls, and that leptin injection into the brain does not suppress dyslipidemia in insulin-deficient RIP-Cre^ΔLEPR^ mice. Leptin administration into the brain combined with acipimox, which lowers blood lipids by suppressing triglyceride lipase activity, can restore normal glycemia in insulin-deficient RIP-Cre^ΔLEPR^ mice, suggesting that excess circulating lipids are a driving-force of hyperglycemia in insulin-deficient RIP-Cre^ΔLEPR^ mice. Collectively, our data demonstrate that LEPRs in RIP-Cre^25Mgn^ neurons significantly contribute to glucose-lowering effects of leptin in an insulin-independent manner by suppression of dyslipidemia.

## Introduction

Central leptin injections can maintain euglycemic ranges without exogenous insulin administration in insulin-deficient rodent models (Fujikawa et al., 2013; Fujikawa et al., 2010; German et al., 2011; Meek et al., 2013; Meek et al., 2018; Xu et al., 2016). Previous studies have unraveled key neuronal components contributing to glucose-lowering effects of central leptin signaling. Leptin receptors (LEPRs) in GABAergic neurons substantially contribute to glucose-lowering effects of leptin in an insulin-independent manner (Fujikawa et al., 2013) and intriguingly leptin-responsive GABAergic neurons are restrictedly positioned to the hypothalamic arcuate nucleus (ARC), dorsomedial nucleus (DMH), and lateral areas (LHA) (Fujikawa et al., 2013; Vong et al., 2011). Of note, among the ARC, DMH, and LHA, the vast of majority of leptin-responsive GABAergic neurons are located in the ARC and DMH (Vong et al., 2011). Recent studies have further shown that LEPRs in agouti-related peptide-expressing neurons (AgRP neurons), which are GABAergic and located in the ARC, are key to glucose-lowering effects of central leptin signaling (Singha et al., 2019; Xu et al., 2018). However, other neuronal groups likely contribute to glucose-lowering effects as well, because intracerebroventricular (i.c.v.) leptin injection still can lower hyperglycemia in insulin-deficient mice lacking LEPRs in AgRP neurons (Singha et al., 2019). Identification of neuronal groups underlying glucose-lowering effects of leptin in an insulin-independent manner has not yet been achieved.

A recent study using single cell RNA-sequence shows that GABAergic neurons in the ARC and median eminence (Arc-ME) complex are composed of distinct genetically-defined neuronal groups (Campbell et al., 2017). AgRP neurons are the most dominant neurons among Arc-ME GABAergic neurons (Campbell et al., 2017). In the same study, the single cell RNA-sequence analysis reveals that neurons expressing Cre recombinase driven by a short fragment of rat insulin promoter transgene (RIP-Cre^25Mgn^ neurons) are distinguished from AgRP neurons, yet, uniquely composed from several neuronal groups (Campbell et al., 2017). A RIP-Cre^25Mgn^ mouse line was originally generated to target pancreatic β-cells (Postic et al., 1999). However, the mice ectopically express Cre recombinase in the central nervous system (CNS) (Song et al., 2010; Wicksteed et al., 2010) due to the nature of genetically-engineering methods in the early era (Harno et al., 2013; Magnuson and Osipovich, 2013). Because RIP-Cre^25Mgn^ mice express Cre recombinase in unique and distinct neurons from conventional hypothalamic neurons such as AgRP neurons (Campbell et al., 2017; Choudhury et al., 2005), the transgenic mice have been utilized for studies investigating the role of hypothalamic neurons that are genetically-uncategorized by conventional genes, in particular studies focusing on the regulation of metabolism (Kong et al., 2012; Ladyman et al., 2017; Wang et al., 2018a; Wang et al., 2014).

RIP-Cre^25Mgn^ neurons regulate energy expenditure through thermogenesis (Kong et al., 2012; Wang et al., 2018a) and glucose metabolism (Ladyman et al., 2017; Wang et al., 2014) in the presence of insulin. Mice lacking LEPRs in RIP-Cre^25Mgn^ cells show aberrant fat metabolism including modest increases in body weight and triglyceride along with augmented circulating insulin (Covey et al., 2006). Because LEPRs are not express in pancreatic β-cells (Fujikawa et al., 2013; Soedling et al., 2015), metabolic phenotypes in mice lacking LEPRs in RIP-Cre^25Mgn^ cells result from ablation of LEPRs in the CNS. Of note, studies show that GABAergic RIP-Cre^25Mgn^ neurons are located in restricted areas, the ARC, DMH, and the medial tuberal nucleus (MTu) (Kong et al., 2012). The anatomical profiling of GABAergic RIP-Cre^25Mgn^ neurons is very similar to leptin-responsive GABAergic neurons, which are positioned in the ARC and DMH (Fujikawa et al., 2013; Vong et al., 2011).

Based on aforementioned studies, we reasoned that LEPRs in RIP-Cre^25Mgn^ neurons significantly contribute to glucose-lowering effects of leptin in an insulin-independent manner. To test our hypothesis, we generated insulin-deficient mice lacking LEPRs in RIP-Cre^25Mgn^ neurons (RIP-Cre^ΔLEPR^) and examined whether i.c.v. leptin injection can lower hyperglycemia in these mice without insulin. Our results indicate that RIP-Cre^25Mgn^ neurons are vital components for glucose-lowering effects of leptin through the regulation of lipid metabolism in an insulin-independent manner.

## MATERIAL AND METHODS

### Genetically-Engineered Mice

RIP-Cre^25Mgn^ (Postic et al., 1999) and Ai9 mice (Madisen et al., 2010) were obtained from the Jackson Laboratory (JAX, USA, #003573 and #007909). *Lepr*^flox/-^ (Balthasar et al., 2004) and *Lepr*^loxTB/WT^ (Berglund et al., 2012) mice were obtained from Dr. Joel Elmquist at the University of Texas Southwestern Medical Center (UTSW) and are also available at the JAX (#008327 and #018989). RIP^Herr^-DTR mice (Thorel et al., 2010) were obtained from Dr. Pedro Herrera at Geneva University. *Gcg*^loxTB/WT^ mice were generated as previously described (Chambers et al., 2017). To generate mice lacking LEPRs in RIP-Cre^25Mgn^ neurons, we bred RIP-Cre^25Mgn^ with *Lepr*^flox/-^ mice followed by breeding RIP-Cre^25Mgn^∷*Lepr*^flox/flox^ with *Lepr*^flox/flox^. To generate mice re-expressing LEPRs in RIP-Cre^25Mgn^ neurons otherwise LEPRs null, we bred RIP-Cre^25Mgn^ with *Lepr*^loxTB/WT^ mice followed by breeding RIP-Cre^25Mgn^∷*Lepr*^loxTB/WT^ with *Lepr*^loxTB/WT^. To pharmacologically induce insulin deficiency, storeptozocin (STZ) was administered (150 mg/kg BW, three times with one week interval) (Fujikawa et al., 2010; Wang et al., 2010). To genetically induce insulin deficiency, RIP^Herr^-DTR mice (Thorel et al., 2010) were bred with mice described above and inject diphtheria toxin. To examine effects of diphtheria toxin injections on the viability of RIP-Cre^25Mgn^ neurons, we introduced the tdTomato allele (Madisen et al., 2010) to identify RIP-Cre^25Mgn^ neurons under the fluorescent microscopy. Mice used are as follows; RIP-Cre^25Mgn^∷*Lepr*^flox/flox^::RIP^Herr^-DTR, *Lepr*^flox/flox^::RIP^Herr^-DTR (control for RIP-Cre^25Mgn^∷*Lepr*^flox/flox^::RIP^Herr^-DTR),RIP-Cre^25Mgn^∷*Lepr*^loxTB/loxTB^::RIP^Herr^-DTR, *Lepr*^loxTB/loxTB^::RIP^Herr^-DTR (null control for RIP-Cre^25Mgn^∷*Lepr*^loxTB/loxTB^::RIP^Herr^-DTR), *Lepr*^WT/WT^::RIP^Herr^-DTR and RIP-Cre^25Mgn^∷*Lepr*^WT/WT^::RIP^Herr^-DTR (wild-type control for RIP-Cre^25Mgn^∷*Lepr*^loxTB/loxTB^::RIP^Herr^-DTR), *Gcg*^loxTB/loxTB^∷RIP^Herr^-DTR, *Gcg*^WT/WT^∷RIP^Herr^-DTR (control for *Gcg*^loxTB/loxTB^ ∷RIP^Herr^-DTR), and RIP-Cre^25Mgn^∷RIP^Herr^-DTR∷Ai9^TB/-^. We used KAPA Mouse genotyping kits (KAPA Biosystems, USA) to determine genotypes. All genotyping primers and predicted band sizes are described in Supplementary Table 1. We used 3-6 month-old male mice whose body weights were above approximately 25 grams. All mice were fed with a normal chow diet (Teklad LM-485, Envigo, US). Animal care was according to established NIH guidelines, and all procedures were approved by the Institutional Animal Care and Use Committee of the University of Texas Southwestern Medical Center and University of Texas Health San Antonio.

### Assessment of basal metabolism prior to induction of insulin deficiency

We measured body weight weekly after weaning at four weeks of age. Blood glucose and plasma insulin were measured at 8-12 weeks of ages prior to inducing insulin deficiency. Body composition of RIP-Cre^25Mgn^∷*Lepr*^loxTB/loxTB^::RIP^Herr^-DTR was measured by rodent fMRI as previously described (Fujikawa et al., 2010; Ramadori et al., 2011; Ramadori et al., 2010). Blood glucose was measured with a commercially available glucose monitor (Bayer Contour, USA). Insulin was measured using a commercially available ELISA kit (Crystal Chem, USA) (RRID:AB_2732074).

### Induction of insulin deficiency by a RIP-DTR approach

To induce insulin deficiency, mice were treated with diphtheria toxin (DT, Sigma, USA). DT was dissolved in sterile 0.9% NaCl solution at a concentration of 150 µg/mL and kept at −80ºC until use. Each concentrated DT aliquot was diluted to 0.075 µg/mL in sterile saline and delivered intraperitoneally (i.p) at a dose of 0.5 µg/kg B.W. one time per day for 3 consecutive days to ablate pancreatic β-cells (Figure 2A). As previously described (Fujikawa et al., 2013; Fujikawa et al., 2010; Singha et al., 2019; Thorel et al., 2010), blood insulin levels were not detectable by the ELISA kit in all mice models except for RIP-Cre^25Mgn^∷*Lepr*^loxTB/loxTB^::RIP^Herr^-DTR and *Lepr*^loxTB/loxTB^::RIP^Herr^-DTR after DT injections (Figure 3H).

**Figure 1.**
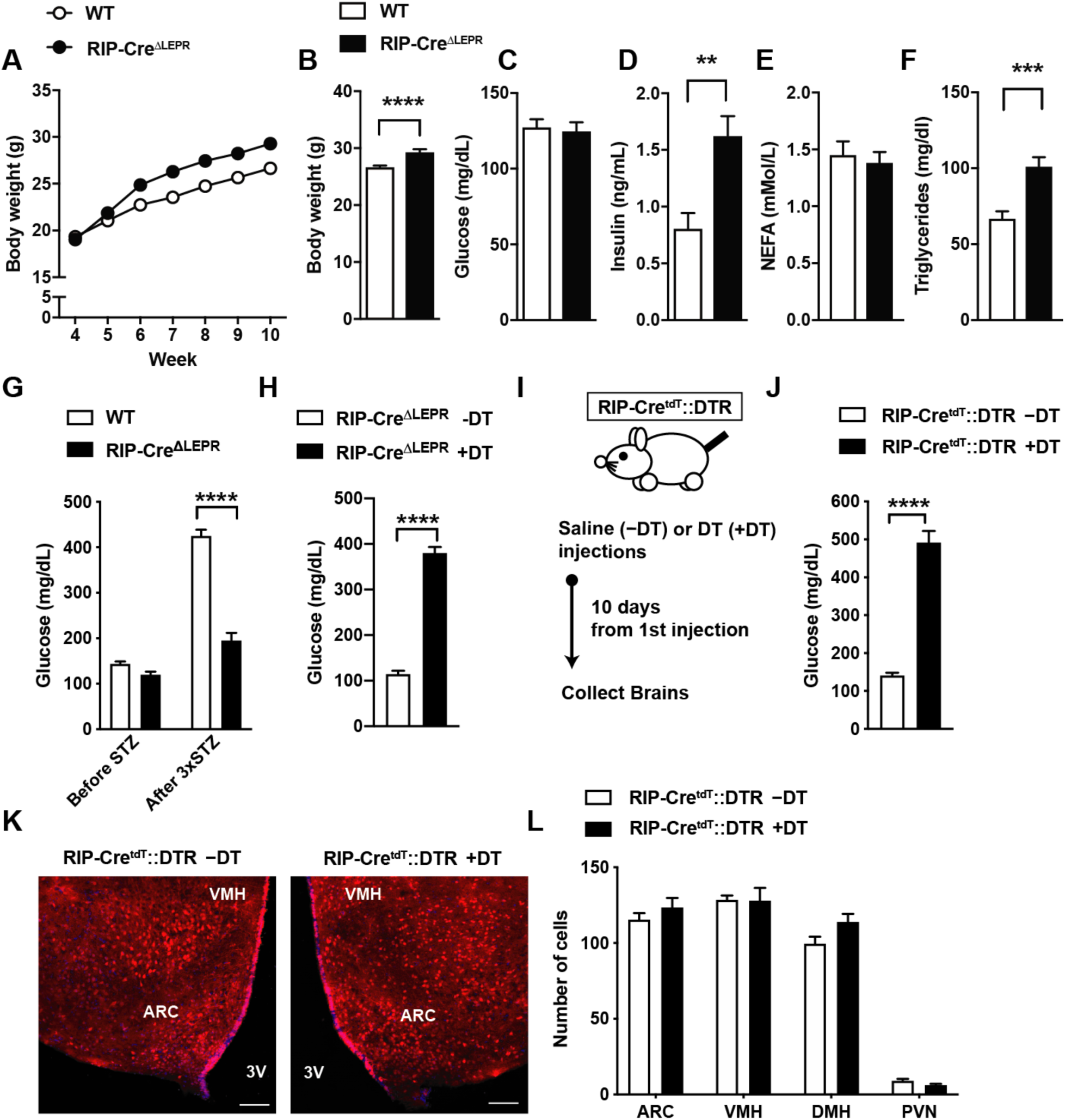
Induction of insulin deficiency with a RIP-DTR approach does not affect RIP-Cre^25Mgn^ neurons. (**A**) The time course of body weight, (**B**) body weight, (**C**) blood glucose, (**D**) plasma insulin, (**E**) plasma NEFA, (**F**) plasma triglyceride in mice lacking LEPRs in RIP-Cre^25Mgn^ neurons (RIP-Cre^ΔLEPR^) at 10 weeks of ages. Wild type control (WT) did not bear RIP-Cre^25Mgn^. Blood glucose levels (**G**) before and 5 days after the final i.p. STZ injection into RIP-Cre^ΔLEPR^ mice. STZ was i.p. administered 3 times (each injection was 5 days apart) at the dose of 150 mg/kg B.W. (**H**) Blood glucose levels 10 days after the induction of insulin deficiency by diphtheria toxin (DT) injections (one injection per day for 3 consecutive days, 0.5 µg/kg B.W.) in RIP-Cre^ΔLEPR^ mice. Control group was administered sterile saline. (**I**) A schematic design for experiments for J, K, and L. Mice were i.p. administered with DT (3 times [0, 1, 2 days], 0.5 µg/kg B.W.) (**J**) Blood glucose levels, (**K**) representative figures (left; Saline injection, right; DT injections, white bar represents 100 µm) and (**L**) the number of tdTomato positive cells in the hypothalamus10 days after the induction of insulin deficiency in RIP-Cre^ΔLEPR^ mice expressing tdTomato in RIP-Cre^25Mgn^ neurons by a RIP-DTR approach. VMH = the ventromedial hypothalamic nucleus, ARC = the hypothalamic arcuate nucleus, 3V, third ventricular, DMH = dorsomedial hypothalamic nucleus, PVN = the paraventricular hypothalamic nucleus. n = 3-33. Values are mean ± S.E.M. **** p < 0.0001, *** p < 0.001, ** p <0.01, * p < 0.05

**Figure 2.**
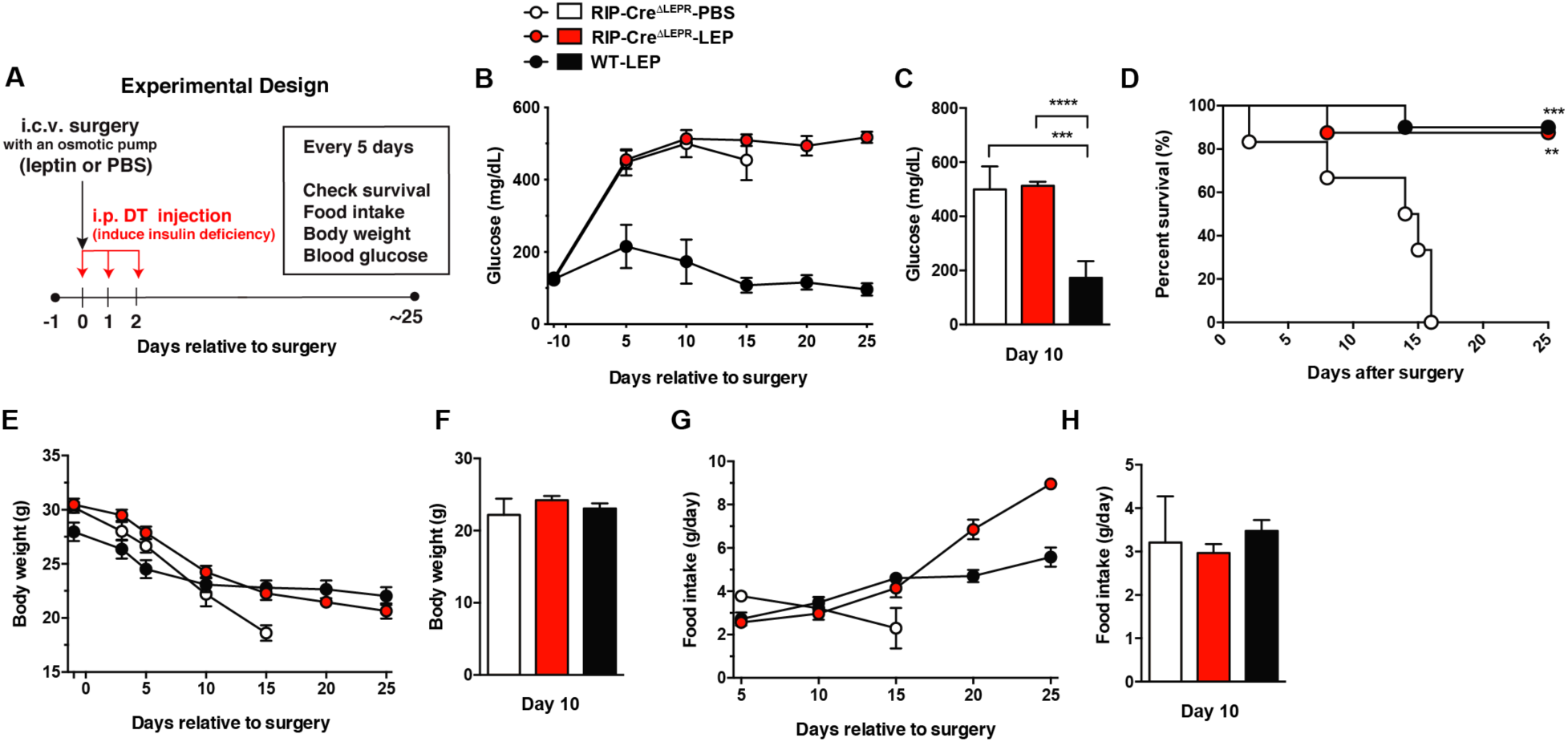
Deletion of LEPRs in RIP-Cre^25Mgn^ neurons blocks glucose-lowering effects of leptin in insulin-deficient mice. (**A**) Experimental design for i.c.v. leptin injection into insulin-deficient mice. (**B**) The time course of blood glucose, (**C**) blood glucose at Days 10, (**D**) the time course of survival percentage, (**E**) the time course of body weight, (**F**) body weight at Days 10, (**G**) the time course of food intake, and (**H**) food intake at Days 10 in insulin-deficient RIP-Cre^ΔLEPR^ mice chronically administered leptin into the lateral ventricle (25 ng/0.11 µL/hour). Insulin deficiency was induced by administration of DT. Control group for leptin administration was administered sterile vehicle (PBS). Genetic control group (WT) did not bear RIP-Cre^25Mgn^. n = 3-8. Values are mean ± S.E.M. **** p < 0.0001, *** p < 0.001, ** p <0.01

**Figure 3.**
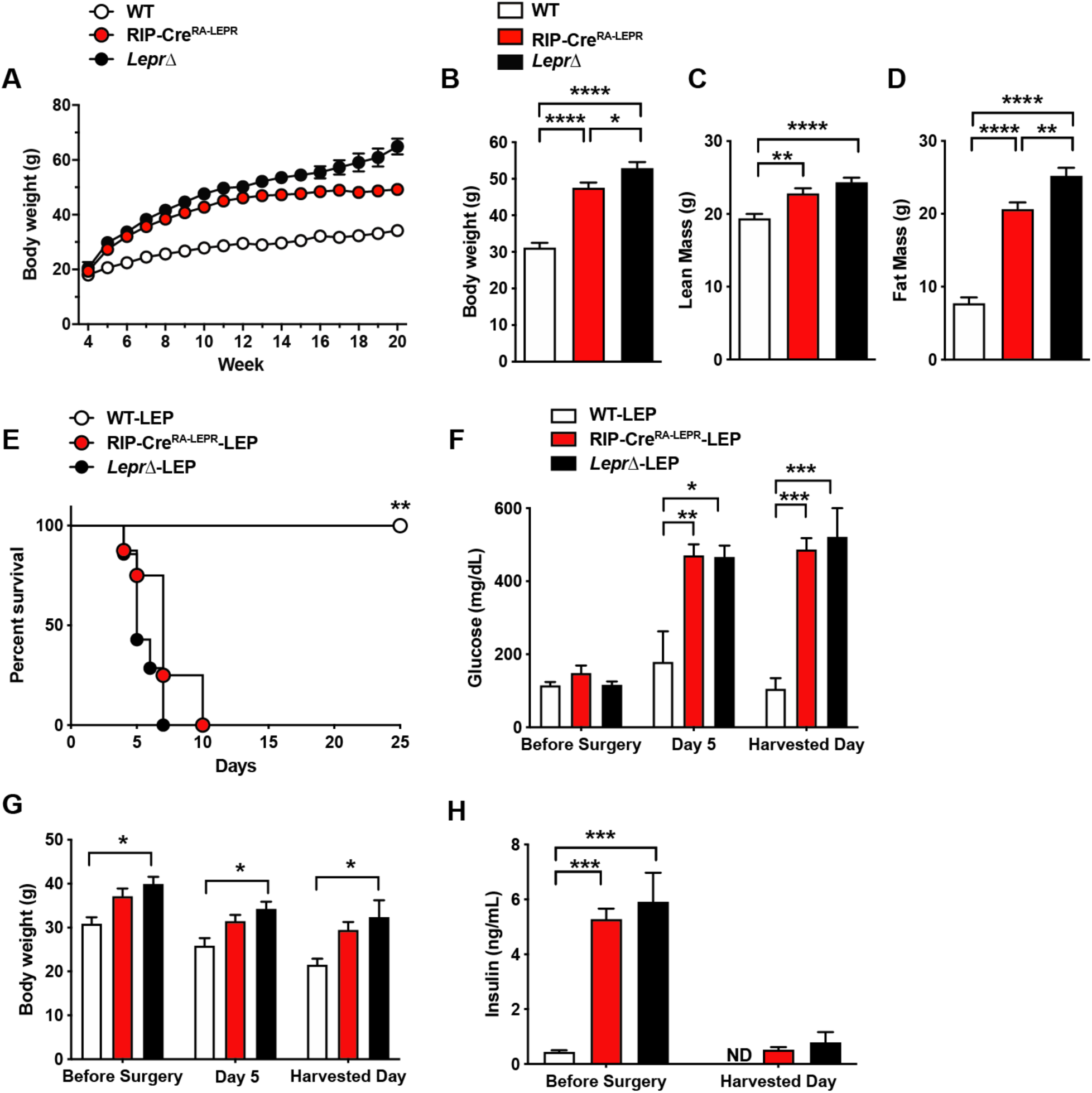
Re-expression of LEPRs in RIP-Cre^25Mgn^ neurons is not sufficient for leptin to lower glucose in insulin-deficient mice. (**A**) The time course of body weight, (**B**) body weight, (**C**) lean mass, and (**D**) fat mass at 15-16 weeks of ages of mice re-expressing LEPRs only in RIP-Cre^25Mgn^ neurons (RIP-Cre^RA-LEPR^). Wild-type control (WT) was composed of RIP-Cre^25Mgn^∷*Lepr*^WT/WT^ and *Lepr*^WT/WT^, and *Lepr*^loxTB/loxTB^ mice were used as LEPRs-deficient mice. (**E**) The survival percentage, (**F**) blood glucose, (**G**) body weight, and (**H**) plasma insulin levels in insulin-deficient RIP-Cre^RA-LEPR^ mice chronically administered leptin into the lateral ventricle (25 ng/0.11 µL/hour). n = 3-14. Values are mean ± S.E.M. **** p < 0.0001, *** p < 0.001, ** p <0.01, * p < 0.05

### Leptin administration into the brain

Leptin (Peprotech, USA; 25 ng/hour/0.11 µL) was dissolved in sterile phosphate-buffered saline (PBS; pH = 7.4, Invitrogen, US) and administered by intracerebroventricular (i.c.v.) infusion using osmotic pumps (Alzet, US) as previously described (Fujikawa et al., 2013; Fujikawa et al., 2010; Singha et al., 2019). An osmotic minipump designed for use in mice (model 1004; Alzet) was implanted subcutaneously and attached via a catheter to the lateral ventricle cannula for i.c.v. administration. PBS was administered to the control group as a placebo treatment. We continuously administered leptin for up to 25 days as pumps are designed to deliver for approximately 28 days.

### Measurement of metabolic parameters and survival

We measured glucose, body weight, and food intake every five days as previously described (Fujikawa et al., 2013; Fujikawa et al., 2010; Singha et al., 2019). We plotted survival to determine if LEPRs in RIP-Cre^25Mgn^ neurons are required or sufficient for leptin’s capacity to reduce lethality in insulin-deficient mice. Free fatty acids, ketone bodies, triglycerides, and glycerol in blood were measured by commercially available kits (Wako diagnose, US; and Cayman US for glycerol) as described previously (Fujikawa et al., 2010).

### Immunohistochemistry

Mice were deeply anesthetized with isoflurane and underwent transcardiac perfusion fixation with 4% paraformaldehyde as previously described (Scott et al., 2009). After cryoprotection in 30% sucrose-sterile-PBS solution, the brain was cut in 25 µm sections using a freezing microtome. Brain sections were mounted on glass slides using the antifade mounting medium with DAPI (H-1500, Vector Lab, USA). Images were captured by fluorescence microscopy (Keyence US, US; Model: BZ-X710). Neurons expressing tdTomato fluorescent and distributed in the hypothalamus at the coronal section approximately −0.5 to −2.0 mm from the caudal to the bregma were manually counted.

### Assessment of mRNA

Mice were deeply anesthetized with isoflurane and tissues were quickly removed, frozen in liquid nitrogen and subsequently stored at –80ºC. RNA was extracted using STAT60 reagent (Amsbio, MA, USA). Complementary DNA from 1 µg of input RNA was generated with the High Capacity cDNA Reverse Transcription Kits (Life Technologies). SYBR Green PCR master mix (Life Technologies) was used for the quantitative real time PCR analysis. Sequences of deoxy-oligonucleotides primers are outlined in supplementary file (Supplemental Table 2).

### Assessment of hepatic lipids

Liver tissue was homogenized in ice-cold diluted phosphate-buffered saline (0.1X PBS) as described previously (Palavicini et al., 2016). Lipids were extracted by a modified procedure of Bligh and Dyer extraction in the presence of internal standards which were added based on the total protein content of individual samples as described previously (Cheng et al., 2006; Cheng et al., 2007; Wang et al., 2018b). A triple-quadrupole mass spectrometer (Thermo Scientific TSQ Altis, CA, USA) and a Quadrupole-Orbitrap™ mass spectrometer (Thermo Q Exactive™, San Jose, CA) equipped with a Nanomate device (Advion Bioscience Ltd., NY, USA) and Xcalibur system software was used as previously described (Han et al., 2008; Wang et al., 2017; Yang et al., 2009). Briefly, diluted lipid extracts were directly infused into the ESI source through a Nanomate device. Typically, signals were averaged over a 1-min period in the profile mode for each full scan MS spectrum. For tandem MS, a collision gas pressure was set at 1.0 mTorr, but the collision energy varied with the classes of lipids. Similarly, a 2- to 5-min period of signal averaging in the profile mode was employed for each tandem MS mass spectrum. All full and tandem MS mass spectra were automatically acquired using a customized sequence subroutine operated under Xcalibur software. Data processing including ion peak selection, baseline correction, data transfer, peak intensity comparison, ^13^C deisotoping, and quantitation were conducted using a custom programmed Microsoft Excel macro as previously described after considering the principles of lipidomics (Wang et al., 2016; Yang et al., 2009).

### Injection of acipimox

Acipimox (Sigma Aldrich, US) was i.p. administered at a dose of 100 mg/kg B.W. two times per day for five consecutive days. Control solution was sterile saline (0.9 % NaCl).

### Data analysis

Data are represented as the group mean ± S.E.M. as indicated in each figure legend. Statistical significance was determined using GraphPad PRISM software (ver8, GraphPad, San Diego, CA) by unpaired t-test, one-way ANOVA followed by Turkey’s multiple comparison test, two-way ANOVA followed by one-way ANOVA (Tukey’s multiple comparison test if the interaction was significant) or unpaired t-test in the same factor (if the interaction was not significant), or repeated measures ANOVA followed by unpaired t-test if the interaction was significant. For analysis of survival curves, Log-rank (Mantel-Cox) testing was used. Since the number of mice surviving declined over time, we were prohibited from utilizing repeated measures ANOVA (Figure 2B, E, and G, and Figure 4D, F, and H). We, therefore, performed statistical analysis and showed each day individually in Fig 2C, F and H and 4E, H, and G. For all tests, statistical significance was set at a critical value of *P*<0.05.

**Figure 4.**
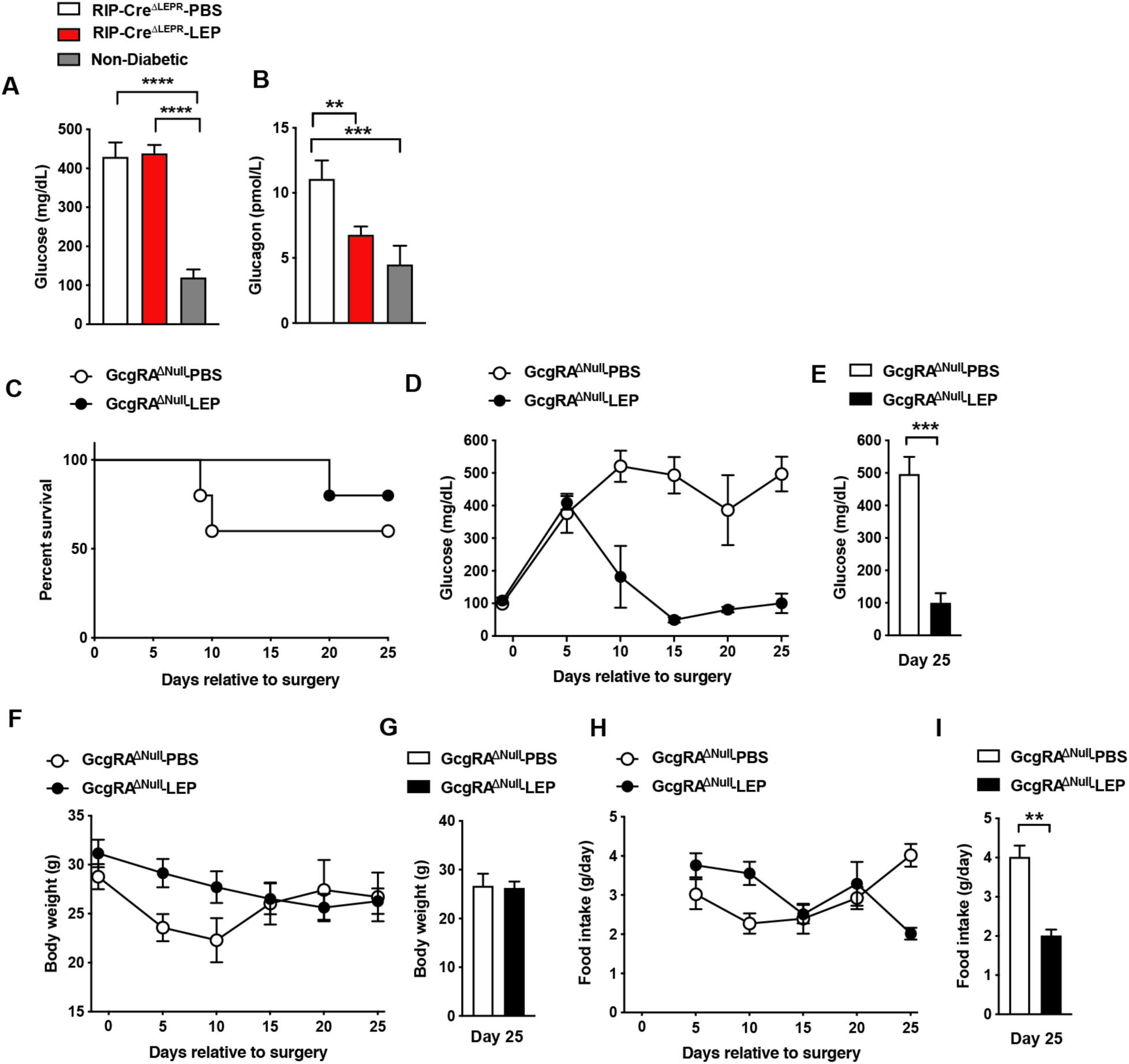
LEPRs in RIP-Cre^25Mgn^ neurons do not contribute leptin-induced suppression of glucagon secretion in insulin-deficient mice. (**A**) Blood glucose and (**B**) plasma glucagon levels in insulin-deficient RIP-Cre^ΔLEPR^ mice 10 days after the induction of chronic administration of leptin into the lateral ventricle (25 ng/0.11 µL/hour). Control group for leptin administration was administered sterile vehicle (PBS). Non-diabetic group was not administered either DT or leptin/PBS. (**C**) The survival percentage, (**D**) the time course of blood glucose, (**E**) blood glucose at Days 25, (**F**) the time course of body weight, (**G**) body weight at Days 25, (**H**) the time course of food intake, and (**I**) food intake at Days 25 in insulin-deficient *Gcg* knockout mice. n = 3-13. Values are mean ± S.E.M. **** p < 0.0001, *** p < 0.001, ** p <0.01.

## Results

### Validation of insulin-deficient RIP-Cre^ΔLEPR^ mice

In agreement with a previous report (Covey et al., 2006), we found that RIP-Cre^ΔLEPR^ mice did not show significant differences of blood glucose and free fatty acid in the presence of insulin, while they exhibited modest obesity and higher circulating insulin and triglyceride levels compared to WT group (Figure 1A-F). To induce insulin deficiency in RIP-Cre^ΔLEPR^ mice, we first used STZ injection methods as we previously described (Fujikawa et al., 2010). However, STZ injections (3 times, 150 mg/kg B.W., each injection was 5 days apart) did not lead to hyperglycemia in RIP-Cre^ΔLEPR^ mice (Figure 1G). Then, we genetically induced insulin deficiency in RIP-Cre^ΔLEPR^ mice with a DT approach (Thorel et al., 2010). To do so, we generate RIP-Cre^ΔLEPR^ mice bearing RIP^Herr^-DTR that induced the expression of DTR in pancreatic β-cells (Thorel et al., 2010). As we previously described (Fujikawa et al., 2013; Thorel et al., 2010), we successfully induced hyperglycemia in RIP-Cre^ΔLEPR^ mice upon DT injections (Figure 1H). As described in the Materials and Methods section, we could not detect plasma insulin in RIP^Herr^-DTR-induced insulin-deficient RIP-Cre^ΔLEPR^ mice (data not shown). Our previous publications show that DT injections into RIP^Herr^-DTR mice did not reduce the number of AgRP and proopiomelanocortin (POMC) neurons, and mRNA levels of *Pomc, Agrp, and Ins2* in the mediobasal hypothalamus. These data suggest that DT injections into RIP^Herr^-DTR mice at the dose we use, which is approximately 100 times less compared to studies ablating neurons (Luquet et al., 2005; Tan et al., 2014), do not ablate hypothalamic neurons. Of note, RIP-Cre^25Mgn^ and RIP^Herr^-DTR were generated by different length of a short fragment of *Ins2* (Herrera, 2000). Additionally, Cre-expression patterns between RIP-Cre^25Mgn^ and RIP-Cre^Herr^ are completely different, for instance, RIP-Cre^Herr^ mice show significantly less Cre activity in the CNS (Rother et al., 2012; Wicksteed et al., 2010). We assumed that DTR is unlikely expressed in all RIP-Cre^25Mgn^ neurons based on previous studies. Nonetheless, we examined if DT injections have deleterious effects on RIP-Cre^25Mgn^ neurons. We administered DT (3 times, 50 ng per kg BW) into RIP-Cre^25Mgn^:: RIP^Herr^-DTR::*tdTomato*^TB/TB^ mice, which allow us to visualize RIP-Cre^25Mgn^ neurons by a red fluorescent reporter (Figure 1I). Ten days after the first injection of DT that was sufficient to induce hyperglycemia (Figure 1J), we examined if DT injections could ablate RIP-Cre^25Mgn^ neurons. We did not find any significant differences of numbers of RIP-Cre^25Mgn^ neurons after DT injections (Figure 1K and L), confirming that our RIP^Herr^-DTR approach does not ablate RIP-Cre^25Mgn^ hypothalamic neurons, similar to AgRP (Singha et al., 2019), POMC and hypothalamic GABAergic neurons (Fujikawa et al., 2013).

### LEPRs in RIP-Cre^25Mgn^ neurons are required, but not sufficient for glucose-lowering effects of leptin

Next, we asked if LEPRs in RIP-Cre^25Mgn^ neurons are required for glucose-lowering effects in an insulin-independent manner. To this end, we administered leptin into the lateral ventricular of insulin-deficient RIP-Cre^ΔLEPR^ mice and examined blood glucose levels in these mice. The experimental design is illustrated in Figure 2A. Intriguingly, chronic i.c.v. leptin injection did not reverse hyperglycemia in insulin-deficient RIP-Cre^ΔLEPR^ mice (Figure 2B and C), suggesting that LEPRs in RIP-Cre^25Mgn^ neurons play a critical role in glucose-lowering effects of leptin in an insulin-independent manner. As we expected, insulin deficiency decreased survival of RIP-Cre^ΔLEPR^ mice administered PBS (RIP-Cre^ΔLEPR^-PBS; Of note, from here, all groups containing either -PBS or -LEP in the abbreviation are insulin-deficient mice) within 2-3 weeks after the induction of insulin deficiency (Figure 2D). Surprisingly, the survival rate of insulin-deficient RIP-Cre^ΔLEPR^ mice administered leptin (RIP-Cre^ΔLEPR^-LEP) was comparable to insulin-deficient WT mice administered leptin (WT-LEP) (Figure 2D). Previously, our studies have shown that there is no correlation between the improvement of blood glucose and survival probability after i.c.v. leptin injection (Fujikawa et al., 2013; Fujikawa et al., 2010; Singha et al., 2019), and this study further confirmed this notion. Body weight between RIP-Cre^ΔLEPR^-LEP and WT-LEP was comparable 10 days after leptin administration was initiated (Figure 2E and F). Food intake was comparable until 15 days after leptin administration was initiated (Figure 2G and H). Previous studies clearly have shown that the amount of food intake could not explain glucose-lowering effects of leptin (Fujikawa et al., 2010; Wang et al., 2010). At Day 10, RIP-Cre^ΔLEPR^-LEP showed significantly higher blood glucose compared to WT-LEP, yet the amount of food intake between RIP-Cre^ΔLEPR^-LEP and WT-LEP was comparable (Figure 2C and H). These data demonstrated that it is unlikely that hyperglycemia in RIP-Cre^ΔLEPR^-LEP resulted from the amount of food intake or body weight differences after induction of insulin-deficiency. However, we cannot exclude the possibility that the increased food intake 20 days after induction of insulin-deficiency contributed to some of the hyperglycemia observed at that time in RIP-Cre^ΔLEPR^-LEP vs. WT-LEP (Figure 2G).

We further asked if expression of LEPRs only in RIP-Cre^25Mgn^ neurons is sufficient for leptin to exert its glucose-lowering effects. To do so, we generated mice re-expressing LEPRs in RIP-Cre^25Mgn^ neurons (RIP-Cre^RA-LEPR^). RIP-Cre^RA-LEPR^ mice had a similar body weight up to 14 to 15 weeks of ages compared to LEPRs null mice (*Lepr*Δ) (Figure 3A). Starting at 15 weeks of age, RIP-Cre^RA-LEPR^ mice had significantly lower body weight along with reductions of fat mass compared to *Lepr*Δ mice (Figure 3B and D). However, the body weight of RIP-Cre^RA-LEPR^ mice was still extremely higher than WT control mice (Figure 3A-D). We chronically administered leptin into the lateral ventricular of DT-injected RIP-Cre^RA-LEPR^mice (RIP-Cre^RA-LEPR^-LEP). We did not see any improvements of the survival rate, blood glucose, body weight of RIP-Cre^RA-LEPR^-LEP compared to *Lepr*Δ-LEP (Figure 2E-H), although these groups still showed a tiny residue of insulin in blood after DT injections. Collectively, these data indicate that LEPRs in RIP-Cre^25Mgn^ neurons are required, but not sufficient for the insulin-independent glucose-lowering effects of leptin.

### Glucose-lowering effects of leptin are independent of glucagon singling

Glucagon is one of the key factors contributing to hyperglycemia in insulin deficiency (Lee et al., 2012; Lee et al., 2011; Unger and Cherrington, 2012). Leptin injection can lower hyperglucagonemia in insulin-deficient rodents (Fujikawa et al., 2010; Wang et al., 2010; Yu et al., 2008), suggesting that suppression of hyperglucagonemia is key for glucose-lowering effects of leptin. We examined blood glucagon levels in RIP-Cre^ΔLEPR^-LEP, however, i.c.v. leptin injection significantly lowered blood glucagon in insulin-deficient RIP-Cre^ΔLEPR^ mice (Figure 4B), while RIP-Cre^ΔLEPR^-LEP showed significantly higher blood glucose levels compared to RIP-Cre^ΔLEPR^-PBS. These data suggest that hyperglucagonemia is not a driving-factor for hyperglycemia seen in RIP-Cre^ΔLEPR^-LEP (Figure 2B and 4A). This result brought the question what a role of glucagon for glucose-lowering effects of leptin is. Intriguingly, previous studies have indicated that glucose-lowering effects of leptin is not necessarily correlated with circulating glucagon levels (German et al., 2010; Perry et al., 2014). We further determined if glucose-lowering effects of leptin can be executed independently of glucagon signaling. To do so, we utilized mice lacking the glucagon gene, *Gcg* (GcgRA^ΔNull^) (Chambers et al., 2017). We chronically i.c.v. administered leptin into insulin-deficient GcgRA^ΔNull^ mice (GcgRA^ΔNull^-LEP) and examined their blood glucose levels, survival rate, body weight, and food intake. I.c.v. PBS injection did not reverse hyperglycemia in insulin-deficient Gcg^KO^ mice (GcgRA^ΔNull^-PBS) (Figure 4D). Interestingly, i.c.v. leptin administration normalized glucose insulin-deficient GcgRA^ΔNull^ mice (Figure 4D and E).

Of note, the percent survival of Gcg^KO^-PBS and were ∼60% at 25 days after induction of insulin deficiency (Figure 4C). All our previous studies have shown that insulin deficiency caused by the RIP^Herr^-DTR method leads to decease in mice within 2-3 weeks (Fujikawa et al., 2013; Singha et al., 2019; Thorel et al., 2010). These facts suggest that hyperglucagonemia has a negative impact on the survivability of insulin-deficient mice independently from its effects on glucose metabolism. We speculated that restoration of glucagon levels by i.c.v. leptin administration (Figure 4B) may contribute to the improvements of survival rate in RIP-Cre^ΔLEPR^-LEP (Figure 2). Further studies will be warranted to investigate the role of glucagon in leptin-induced reverse effects of lethality. Collectively, these data indicate that glucose-lowering effects of leptin in an insulin-independent manner unlikely rely on glucagon system.

### LEPRs in RIP-Cre^25Mgn^ neurons mediate anti-dyslipidemia effects of leptin in an insulin-independent manner

Leptin can improve dyslipidemia in insulin-deficient rodents (Fujikawa et al., 2010; Perry et al., 2014; Wang et al., 2010). Recent studies have pinpointed that improvements of aberrant fat metabolism contributes to glucose-lowering effects of leptin in insulin-deficient rodents (Denroche et al., 2015; Perry et al., 2017; Perry et al., 2014). For instance, administration of fatty acid emulsion into bloodstream (Perry et al., 2014) or i.p. glycerol injection (Denroche et al., 2015) can reverse glucose-lowering effects of leptin in insulin-deficient rodents. Excess lipids in blood disrupts hepatic glucose metabolism, leading to excess hepatic glucose production (Perry et al., 2017). To determine whether dyslipidemia could contribute to preventing leptin for glucose-lowering effects in insulin-deficient RIP-Cre^ΔLEPR^ mice, we first measured circulating free fatty acids (FFAs), ketone bodies, glycerol, and triglyceride in RIP-Cre^ΔLEPR^-LEP at 10 days after the beginning of leptin administration. Intriguingly, RIP-Cre^ΔLEPR^-LEP and RIP-Cre^ΔLEPR^-PBS showed significantly higher levels of all of these fat metabolism substrates (Figure 5A-D). In particular, blood triglyceride levels in RIP-Cre^ΔLEPR^-LEP and RIP-Cre^ΔLEPR^-PBS were significantly higher than WT-LEP. We further examined hepatic free fatty acid and triglyceride levels. Hepatic FFAs and triglyceride levels in RIP-Cre^ΔLEPR^-LEP and RIP-Cre^ΔLEPR^-PBS were extremely higher compared to WT-LEP and WT-PBS (Figure 5E and F). To determine if liver fat synthesis is dysregulated that leads to the high levels of FFAs and triglyceride in RIP-Cre^ΔLEPR^-LEP and RIP-Cre^ΔLEPR^-PBS, we measured mRNA levels of genes related to fatty acid synthesis and oxidation. As previously reported (Wang et al., 2010), leptin can restore mRNA levels of *Srebp-1c* and *Scd-1* in liver (Figure 5G) in an insulin-independent manner. Interestingly, i.c.v. leptin administration did not restore mRNA levels of *Srebp-1c* and *Scd-1* in liver of insulin deficient RIP-Cre^ΔLEPR^ mice (Figure 5G). We assume that excess hepatic lipids suppress genes related to *de novo* lipid synthesis (Horton et al., 2002) in RIP-Cre^ΔLEPR^-LEP and RIP-Cre^ΔLEPR^-PBS. These data indicate that liver in RIP-Cre^ΔLEPR^-LEP and RIP-Cre^ΔLEPR^-PBS unlikely generated an excess amount of *de novo* lipids, and we asked if lipids synthesis in adipose tissues may increase by assessing mRNA of genes related to lipogenesis and lipolysis. Central leptin signaling suppresses lipogenesis in white adipose tissues (WAT) of mice in the insulin-clamped condition (Buettner et al., 2008). Because insulin deficiency induces the drastic reduction of WAT weight due to augmented lipolysis and decreased lipogenesis, we had difficulties in collecting WAT of mice in all groups at Days 10 (Figure 2 and Supplemental Figure 1B), therefore, we collected WAT at Days 5. We did not observe drastic increases in mRNA levels of gene related to lipogenesis and lipolysis in perigonadal WAT of RIP-Cre^ΔLEPR^-LEP and RIP-Cre^ΔLEPR^-PBS (Figure 5H). Rather, we found that mRNA levels of *Fasn, Pnpla*, and *Lipa* in RIP-Cre^ΔLEPR^-LEP were significantly lower compared to WT-PBS, suggesting that the excess blood lipids did not resulted from *de novo* synthesis of lipid or lipolysis. Because the amount of food intake in RIP-Cre^ΔLEPR^-LEP was comparable of that in WT-LEP at 5 days and 10 days (Figure 2 and Supplemental Figure 1C) and we used normal chow diet that contains relatively low fat (17 % of total calories), excess lipids unlikely result from dietary lipids. Our previous studies indicate that i.c.v. leptin injection restores glucose utilization in skeletal muscle of insulin-deficient mice, and mRNA levels of genes related to glucose oxidative pathway suggest that leptin can also restore glucose utilization in skeletal muscle of RIP-Cre^ΔLEPR^ mice (Supplemental Figure 1D). Further studies will be warranted to identify organs/cells that generate excess circulating lipids in insulin-deficient RIP-Cre^ΔLEPR^ mice.

**Figure 5.**
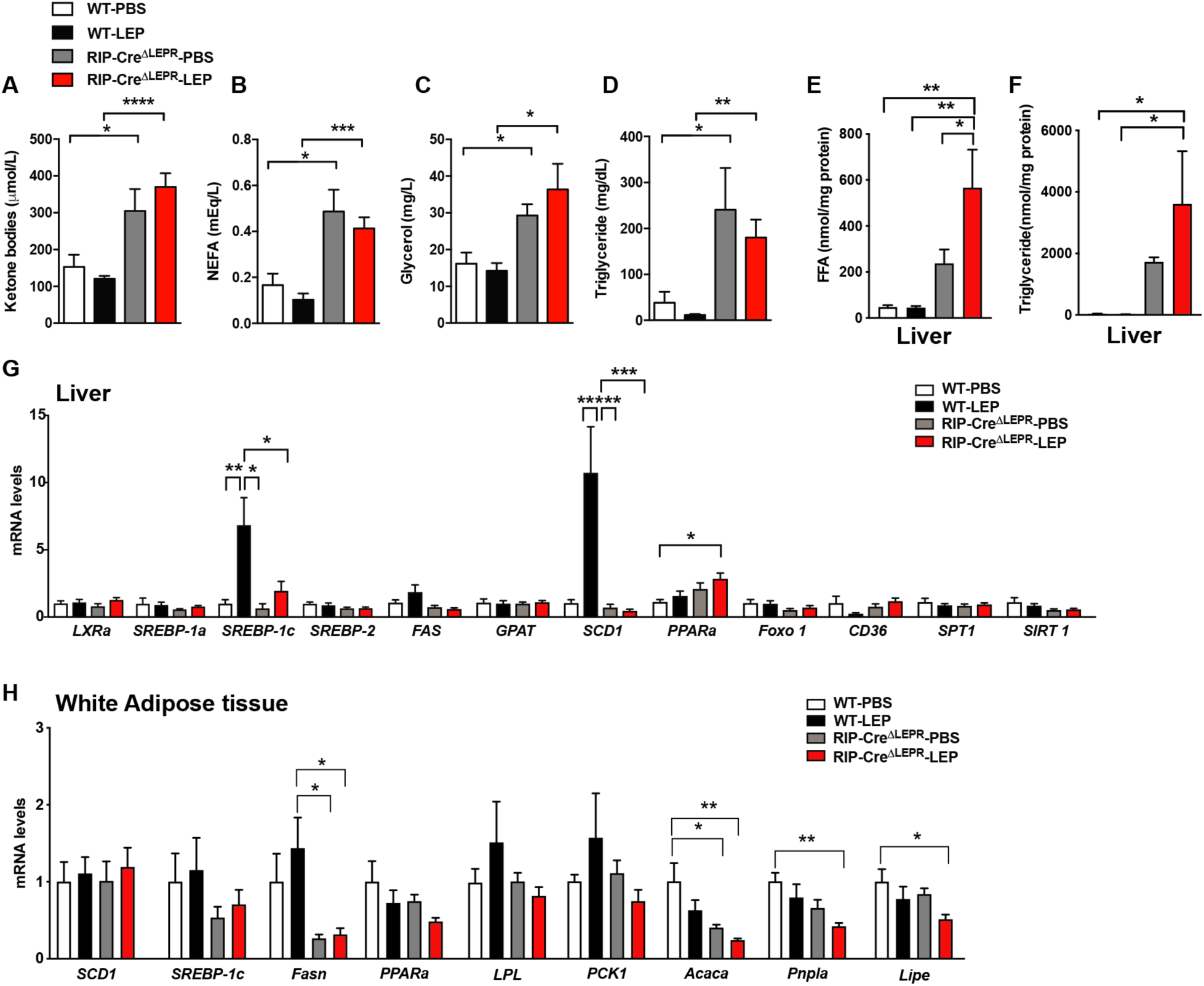
LEPRs in RIP-Cre^25Mgn^ neurons mediate anti-dyslipidemia effects of leptin in insulin-deficient mice. (**A**) Plasma ketone bodies, (**B**) non-esterified fatty acids (NEFA), (**C**) Glycerol, (**D**) Triglyceride, (**E**) hepatic total free fatty acids level, (**F**) hepatic total triglyceride levels, mRNA levels of genes related to lipid metabolism in (**G**) liver and (**H**) perigonadal white adipose tissue of insulin-deficient RIP-Cre^ΔLEPR^ mice administered leptin into the lateral ventricle (25 ng/0.11 µL/hour). Blood and tissue samples were collected 5 days (**H**) and 10 days (**A** to **G**) after the induction of chronic administration of leptin into the lateral ventricle. Control groups were administered sterile PBS. n = 4-9. Values are mean ± S.E.M. **** p < 0.0001, *** p < 0.001, ** p <0.01, * p < 0.05

Finally, we asked if suppression of excess blood lipids by acipimox, which reduces fatty acid and triglyceride in blood (Lee et al., 1996), can reverse hyperglycemia in RIP-Cre^ΔLEPR^-LEP. An acipimox injection alone (i.p. 100 mg/kg BW) dramatically reduced blood free fatty acids but not glucose levels in RIP^Herr^-DTR-induced insulin-deficient mice (Figure 6A and B). I.p. administration acipimox (100 mg/kg BW) into RIP-Cre^ΔLEPR^-LEP (two times per day for 5 days) 10 days after leptin administration initiated (Figure 6C) significantly improved hyperglycemia RIP-Cre^ΔLEPR^-LEP (RIP-Cre^ΔLEPR^-LEP-Acip) compared to the control group (RIP-Cre^ΔLEPR^-LEP-PBS) (Figure 6D). Collectively, our studies indicate that lipid-lowering actions are critical for glucose-lowering effects of leptin and LEPRs in RIP-Cre^25Mgn^ neurons mediate anti-dyslipidemia actions of leptin in an insulin-independent manner.

**Figure 6.**
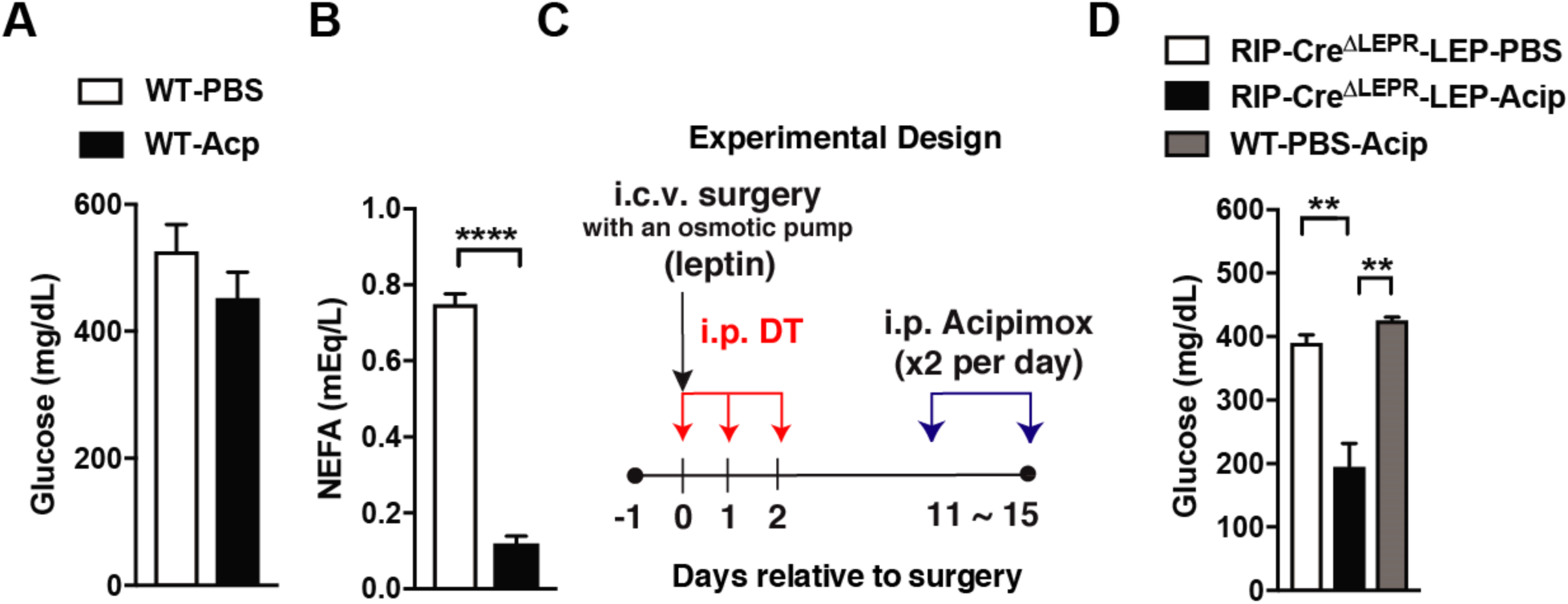
Lowering blood lipids levels reverses hyperglycemia in insulin-deficient RIP-Cre^ΔLEPR^ mice administered leptin. (**A**) Blood glucose and (**B**) plasma NEFA levels in insulin-deficient mice administered Acipimox (WT-Acip) or vehicle (PBS, WT-PBS). RIP-DTR mice were administered DT as described in Figure 1 to induce insulin deficiency. (**C**) Experimental design for D. (**D**) Blood glucose level 5 days after i.p. acipimox injection (2 times per day for 5 days at the dose of 100 mg/kg B.W.) into insulin-deficient RIP-Cre^ΔLEPR^ mice chronically administered leptin into the lateral ventricle (25 ng/0.11 µL/hour) (RIP-Cre^ΔLEPR^-LEP-Acip). Control group for i.p. acipimox injection was administered i.p. saline into insulin-deficient RIP-Cre^ΔLEPR^ mice chronically administered leptin into the lateral ventricle RIP-Cre^ΔLEPR^-LEP-Sal). Control group to determine effects of chronic acipimox injection itself on glucose levels in insulin-deficient mice was WT mice administered i.c.v. PBS and i.p. acipimox (WT-PBS-Acip). n = 4-9. Values are mean ± S.E.M. **** p < 0.0001, *** p < 0.001, ** p <0.01, * p < 0.05

## Discussion

In the present study, we identify that LEPRs in RIP-Cre^25Mgn^ neurons are required for glucose-lowering effects of leptin in an insulin-independent manner as i.c.v. leptin administration does not restore normal glycemia in insulin-deficient mice lacking LEPRs in RIP-Cre^25Mgn^ neurons. Our data suggest that glucagon signaling does not drive hyperglycemia in insulin-deficient mice lacking LEPRs in RIP-Cre^25Mgn^ administered i.c.v. leptin (Figure 4). Rather, our data indicate that excess circulating lipids contribute to the refractory responses of insulin-deficient mice lacking LEPRs in RIP-Cre^25Mgn^ neurons to glucose-lowering effects of leptin in an insulin-independent manner (Figure 5 and 6). Collectively, we propose that LEPRs in RIP-Cre^25Mgn^ neurons are key to regulate lipids metabolism in an insulin-independent manner, although target peripheral tissues have to be determined in future studies.

Our approaches in this study could not allow us to decipher the precise anatomical location of RIP-Cre^25Mgn^ neurons contributing to the regulation of lipid metabolism because of the broad expression pattern of Cre recombinase in RIP-Cre^25Mgn^ mice (Song et al., 2010; Wicksteed et al., 2010) and that LEPRs are also expressed broadly throughout the hypothalamus (Elmquist et al., 1998; Scott et al., 2009). Nonetheless, we assume that GABAergic RIP-Cre^25Mgn^ neurons in the ARC and/or DMH are key for anti-dyslipidemia actions of leptin in insulin-deficient mice, because (1) leptin-responsive GABAergic neurons are located only in the ARC, DMH, LHA, and (2) GABAergic RIP-Cre^25Mgn^ neurons are anatomically limited to the ARC, DMH, and MTu; therefore, ARC and DHM are only overlapped regions that match to the anatomical and chemical-classification profiling from previous studies. Although studies has shown that GABAergic ARC (e.g., AgRP neurons) (Aponte et al., 2011; Betley et al., 2013; Krashes et al., 2011; Steculorum et al., 2016) and DMH (e.g., LEPRs-neurons) (Garfield et al., 2016) neurons regulate multiple aspects of metabolism including food intake and energy expenditure, the role of these neurons in the regulation of lipids metabolism, in particular in an insulin-independent manner, remains unclear.

Interestingly, RIP-Cre^25Mgn^ neurons in the ARC (ARC RIP-Cre^25Mgn^ neurons) are distinct from AgRP neurons (Campbell et al., 2017), which also contribute to glucose-lowering effects of leptin in an insulin-independent manner (Singha et al., 2019; Xu et al., 2018). ARC RIP-Cre^25Mgn^ neurons are composed of at least 9 genetically distinguished neuronal groups (Campbell et al., 2017). A previous study shows that leptin acts on ARC GABAergic RIP-Cre^25Mgn^ neurons to increase energy expenditure along with thermogenesis in interscapular brown adipose tissue without affecting food intake behavior (Kong et al., 2012). Mice lacking LEPRs in RIP-Cre^25Mgn^ neurons exhibit lower energy expenditure without changing food intake compared to control mice (Covey et al., 2006). Compared to ARC RIP-Cre^25Mgn^ neurons, the genetic property of RIP-Cre^25Mgn^ neurons in the DMH (DMH RIP-Cre^25Mgn^ neurons) at the single cell level is still undetermined. In addition, the role of DMH RIP-Cre^25Mgn^ neurons in the regulation of metabolism is completely unknown. Further studies will be warranted to pinpoint the specific neuronal group(s) within RIP-Cre^25Mgn^ neurons that regulate fat metabolism in an insulin-independent manner.

Glucose-lowering effects of leptin are abolished by lipids infusion into insulin-deficient rodents (Perry et al., 2014). Leptin is well known to suppress the hypothalamus-pituitary gland-adrenal gland (HPA)-axis activity including reducing blood corticosterone levels (Ahima et al., 1996). Perry and her colleagues propose that anti-dyslipidemia actions of leptin is mediated by the HPA-axis because infusion of corticosterone reverses leptin-induced improvement effects on aberrant blood free fatty acids and ketone bodies in STZ-administered rats (Perry et al., 2014). Of note, Morton and his colleagues argue that HPA-axis does not contribute to glucose-lowering effects of leptin because corticosterone administration in adrenalectomized rats does not reverse the glucose-lowering effects of leptin (Morton et al., 2015). GABAergic RIP-Cre^25Mgn^ neurons (Kong et al., 2012) and LEPRs-expressing neurons (Elmquist et al., 1998; Scott et al., 2009) do not exist in the hypothalamic paraventricular nucleus; the region of the CNS that releases corticotrophin releasing factor, suggesting that the HPA-axis may not directly play a role in our results. We did not assess if LEPRs in RIP-Cre^25Mgn^ neurons regulate HPA axis, and further investigation will be needed to unravel whether the mechanism by which LEPRs in RIP-Cre^25Mgn^ neurons regulate lipids metabolism in an insulin-independent manner is mediated by HPA-axis.

In summary, our current study demonstrates that LEPRs in RIP-Cre^25Mgn^ neurons significantly contribute to glucose-lowering effects of leptin in an insulin-independent manner by restoring aberrant lipid metabolism. It is still unclear whether LEPRs in RIP-Cre^25Mgn^ neurons mediate effects of leptin on lipid metabolism such as increases of fatty acid oxidation and lipolysis in adipose tissues in the presence of insulin. Unraveling the mechanism by which LEPRs in RIP-Cre^25Mgn^ neurons regulate lipid metabolism may pave a way to design new treatments for several forms of diabetes.

## Acknowledgement

We would like to thank Nancy Gonzalez for the technical assistant. This work was supported by the University of Texas System (UT Rising STARs to T.F.), the American Heart Association (Scientist Development Grant 14SDG17950008 to T.F.), and the National Institutes of Health (1RF1AG061872 to X.H.).

## Contribution

A.K.S. performed and analyzed experiments, and edited the manuscript. J.P.P designed, performed and analyzed experiments, and edited the manuscript. M.P. performed and analyze experiments. D. S. generated *Gcg*^loxTB/WT^ mice and edited manuscript. X.H. supervised experiments and edited manuscript. T.F. designed, performed, supervised, and analyzed experiments, and wrote and finalized the manuscript.

## Figure legends

**Supplemental Figure 1.**
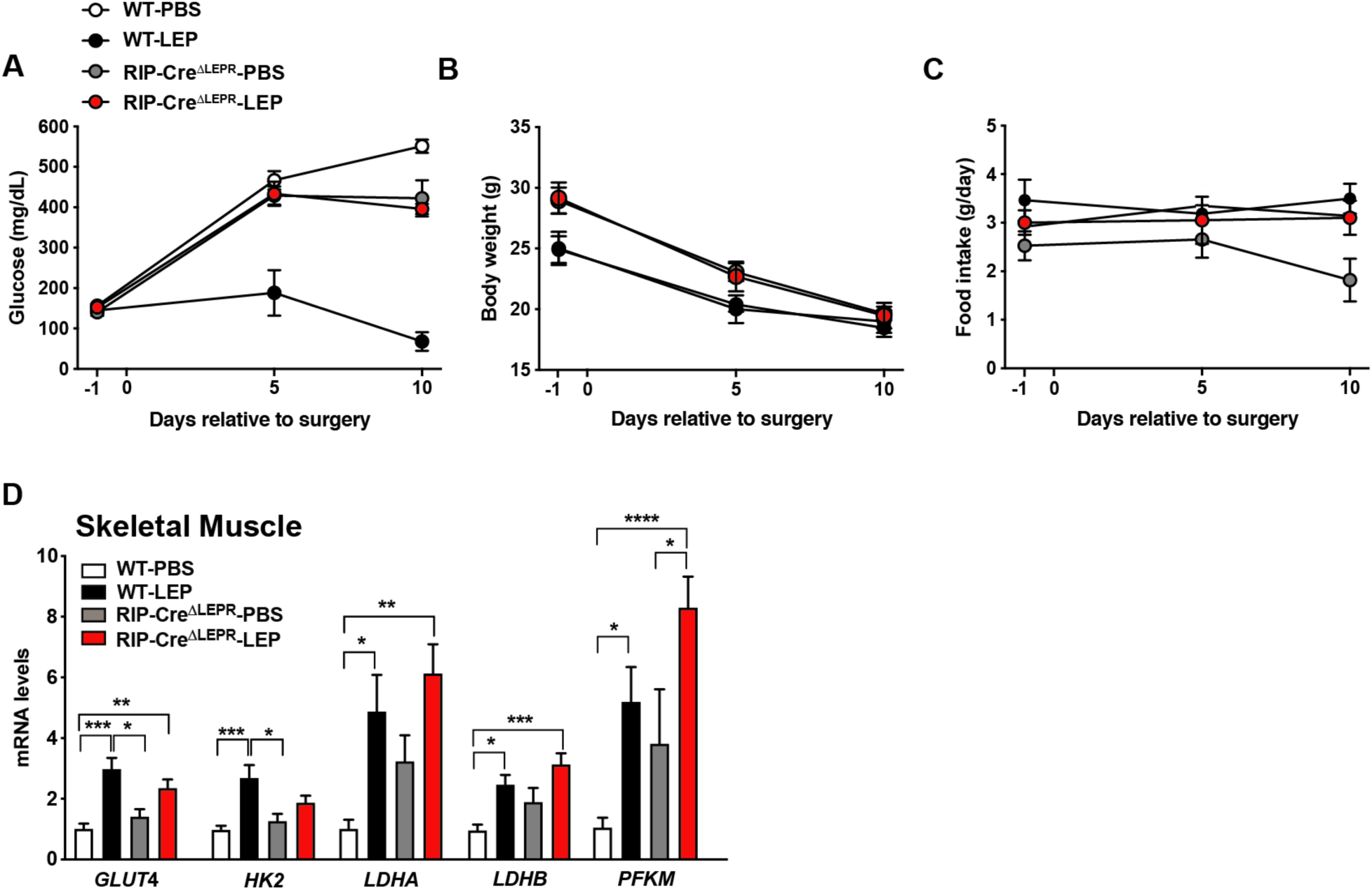
Related to Figure 5. Deletion of LEPRs in RIP-Cre^25Mgn^ neurons unlikely affect mRNA levels of genes related to glucose metabolism in skeletal muscle. (**A**) The time course of glucose, (**B**) body weight, and (**C**) food intake in insulin-deficient RIP-Cre^ΔLEPR^ mice chronically administered leptin into the lateral ventricle (25 ng/0.11 µL/hour). (**D**) mRNA levels of genes in skeletal muscle 10 days after the induction of chronic administration of leptin into the lateral ventricle. Insulin deficiency was induced by administration of DT. Control group for leptin administration was fadministered sterile vehicle (PBS). Genetic control group (WT) did not bear RIP-Cre^25Mgn^. n = 4-9. Values are mean ± S.E.M. **** p < 0.0001, *** p < 0.001, ** p <0.01

**S Table 1.**
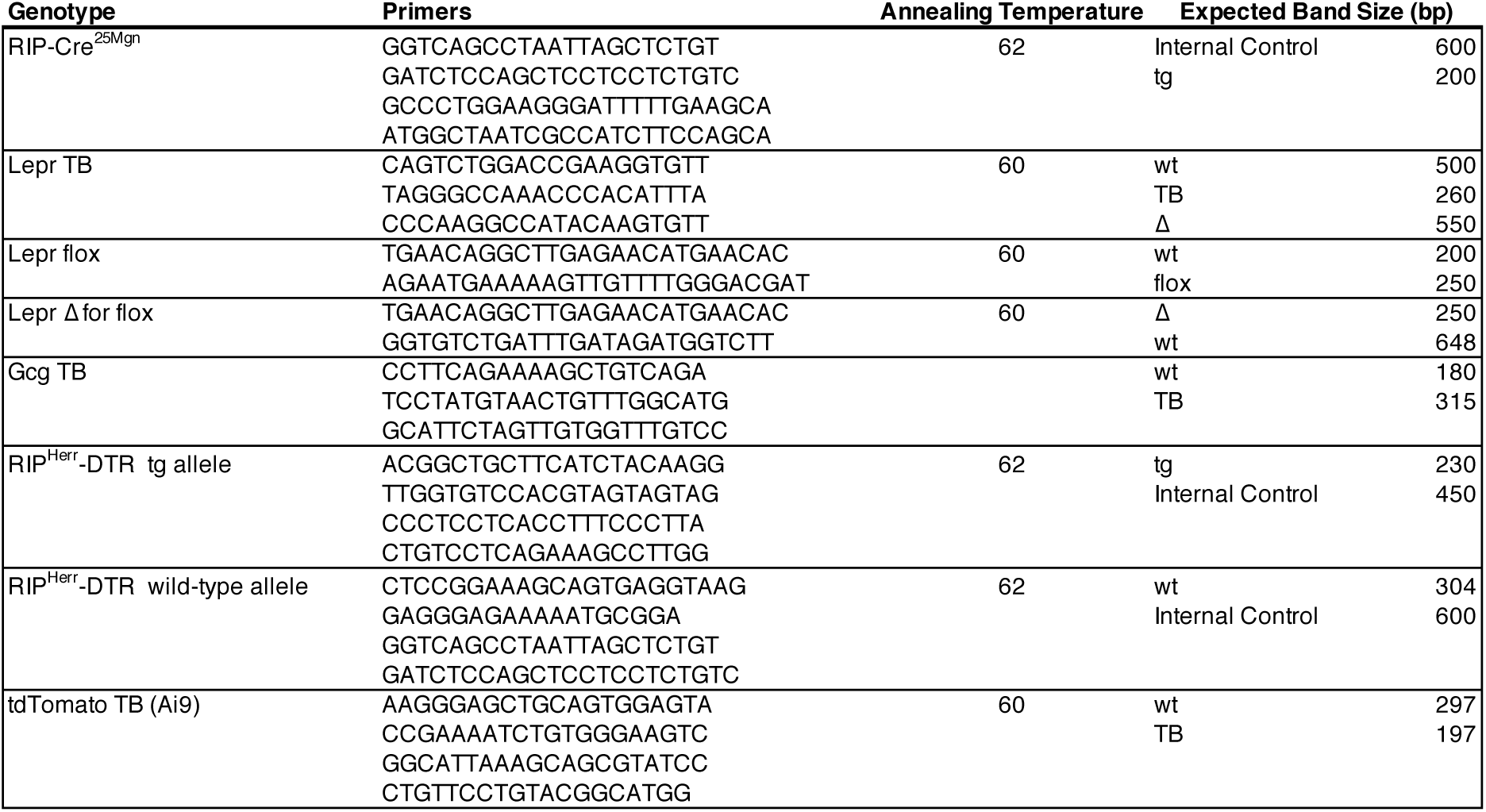
Sequences of genotyping primers.

**S Table 2.**
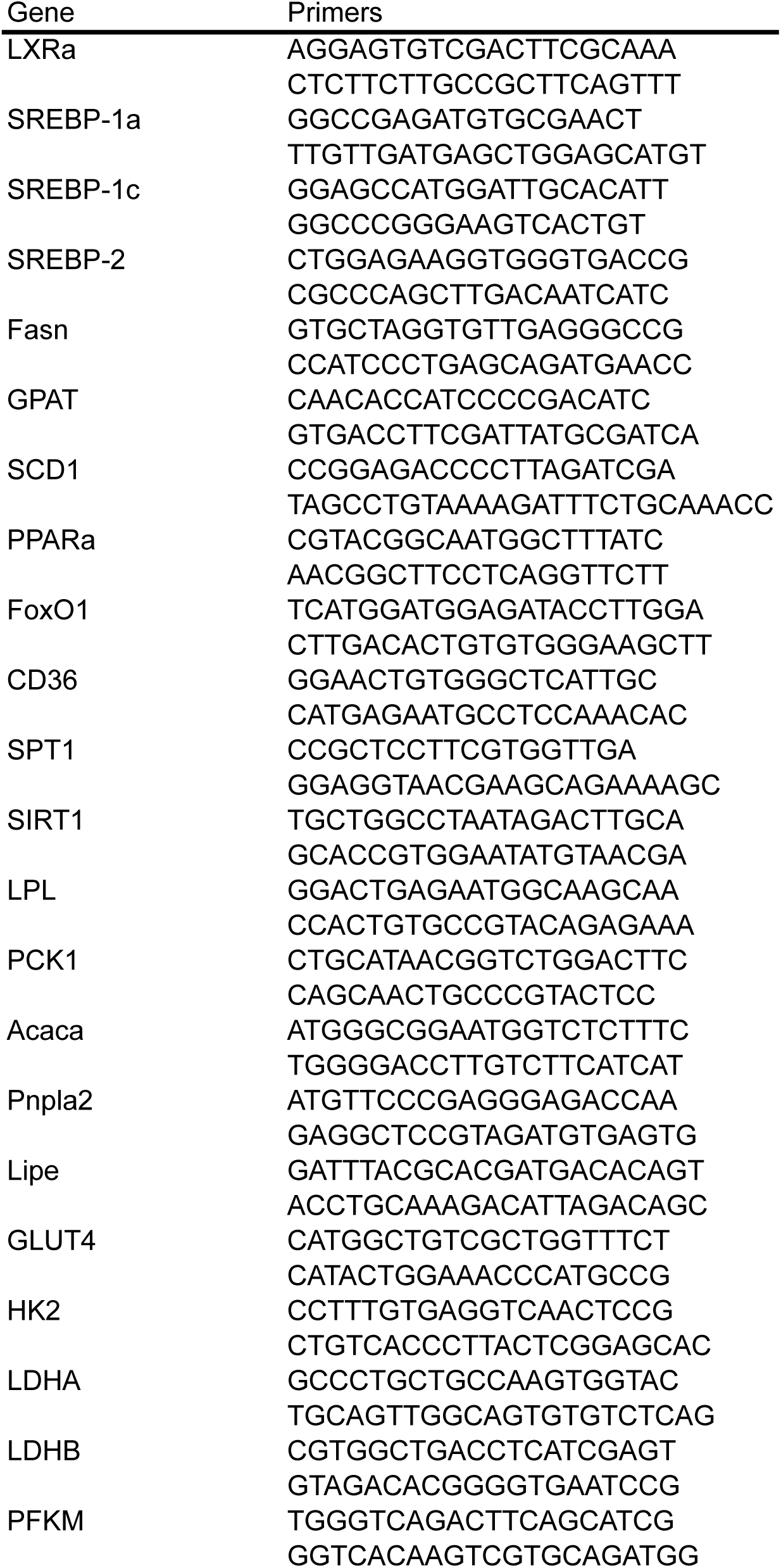
Sequences of qPCR primres.

